# BOFdat: generating biomass objective function stoichiometric coefficients from experimental data

**DOI:** 10.1101/243881

**Authors:** Jean-Christophe Lachance, Jonathan M. Monk, Colton J. Lloyd, Yara Seif, Bernhard O. Palsson, Sébastien Rodrigue, Adam M. Feist, Zachary A. King, Pierre-Étienne Jacques

## Abstract

Genome-scale models (GEMs) rely on a biomass objective function (BOF) to predict phenotype from genotype. Here we present BOFdat, a Python package that offers functions to generate biomass objective function stoichiometric coefficients (BOFsc) from macromolecular cell composition and relative abundances of macromolecules obtained from omic datasets. Growth-associated and non-growth associated maintenance (GAM and NGAM) costs can also be calculated by BOFdat.

BOFdat is freely available on the Python Package Index (pip install BOFdat). The source code and an example usage (Jupyter Notebook and example files) are available on GitHub (https://github.com/jclachance/BOFdat). The documentation and API are available through ReadTheDocs (https://bofdat.readthedocs.io).

## 1 Introduction

Genome-scale models (GEMs) can be used to predict achievable physio-logical states given the constraints imposed by a metabolic network (O’Brien et al., 2015). When the GEM’s objective is to maximize growth, the biomass objective function (BOF) is used to represent a reaction comprising all constituents of the organism (Feist and Palsson, 2010). The metabolites present in the BOF and their coefficients should ideally be obtained from experimental measurements performed in the desired conditions in order to better reflect the organism’s growth requirements (Feist and Palsson, 2010; O’Brien et al., 2015). Despite its importance, the BOF is often mistreated when building GEMs (Xavier et al., 2017; Chan et al., 2017). The widely used Constraint-Based Recon-struction and Analysis toolbox for building metabolic models in Python, COBRApy (Ebrahim et al., 2013), does not include any functionality to incorporate experimental data into a BOF. Here we introduce BOFdat, a COBRApy compatible Python package that facilitates the integration of experimental data for the determination of the BOF.

## 2 Implementation and features

Biomass objective function stoichiometric coefficients (BOFsc) for five macromolecular categories (DNA, RNA, proteins, lipids and metab-olites) can be generated with BOFdat. Accurate calculation of BOFsc by BOFdat is performed as described in Thiele and Palsson, 2010 (Thiele and Palsson, 2010) using both the macromolecular weight fractions and the relative abundance of metabolites from omic datasets (Fig. 1). The architecture of the package is designed so that the BOFsc for each macromolecular category is generated independently using the appropriate `generate_coefficients()` function (Fig. S1). The resulting dictionary is then used with the corresponding `up-date_biomass()` function to update the BOF of the model. The parameters for these functions – thoroughly described in the documentation (bofdat.readthedocs.io) – are mainly the sequence file of the organism, the macromolecular weight fraction of the category, and a comma-separated file containing the relative abundance of each molecule composing that category (Fig. 1). For metabolites and lipids, functions are provided to compare the experimental data with the prediction from the GEM, which allows informed decisions about compounds to include in the BOF (Fig. S1). Finally, a function to calculate ATP maintenance cost is also provided in BOFdat, using input growth data from different carbon sources, along with uptake and secretion rates (Fig. S1).

**Fig. 1 -.**
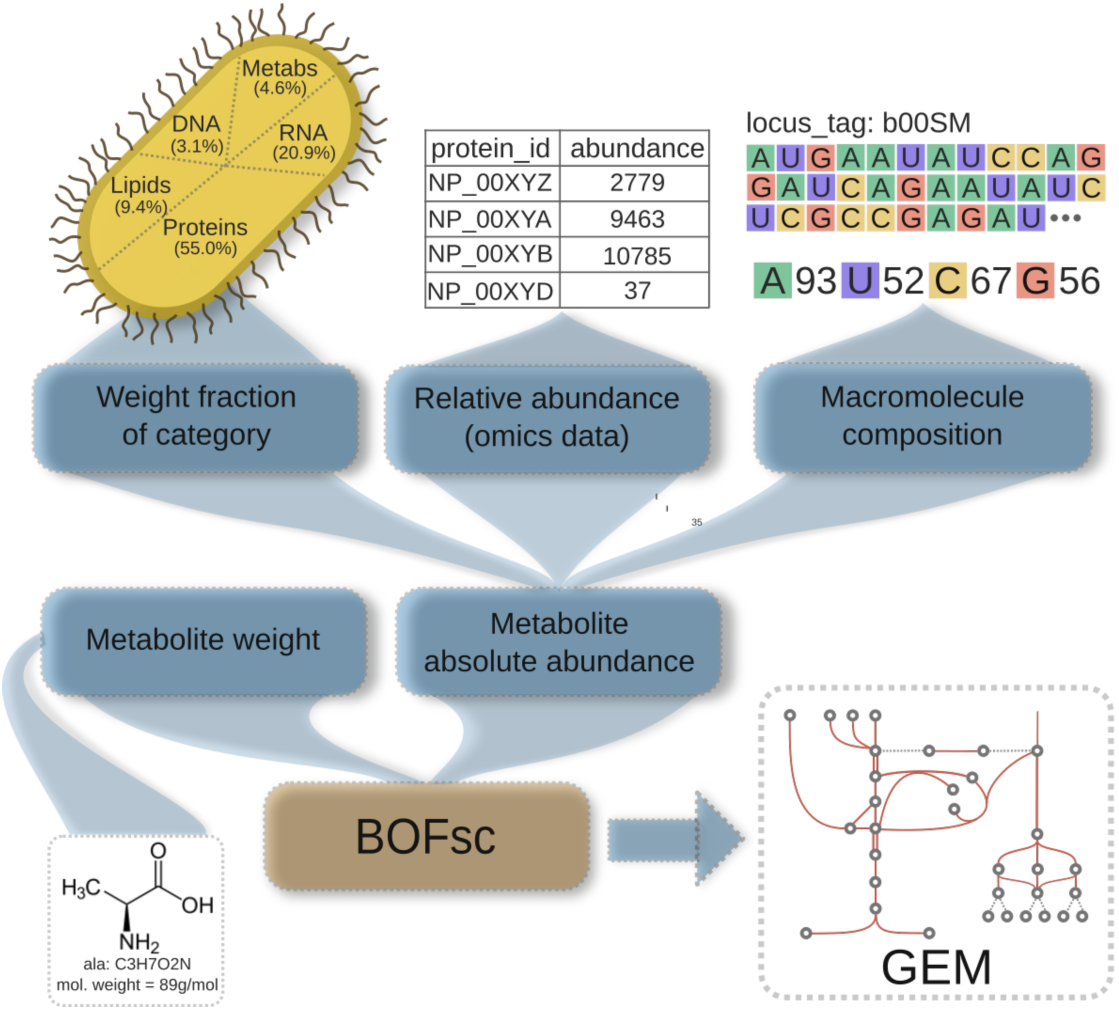
BOFdat overview. Metabolite absolute abundance is calculated based on the integration of the macromolecular category weight fraction (e.g. total protein weight per total cell weight), the relative abundance of each element composing a category (omics data), and the macromolecule composition (sequence). The weight and absolute abundance of each metabolite are in turn used to calculate BOFsc and to update the GEM.

## 3 Usage and application

To validate the use of BOFdat, available experimental omic datasets (Hayashi et al., 2006; Seo et al., 2014; Schmidt et al., 2016; Cheng et al., 2014; Oursel et al., 2007) were used to reconstruct the BOFsc of the high-quality iML1515 model of *Escherichia coli* K-12 MG1655 model (Monk et al., 2017; King et al., 2016), (see the example usage github.com/jclachance/BOFdat). BOFdat was able to generate BOFsc highly similar to those originally included in iML1515 (Fig. S2) based on experimental data only. When varying the weight fraction for each macromolecular category, the minimal error was found within 2% of the experimental weight fraction from the iML1515 BOFsc (Neidhardt et al., 1996; Feist et al., 2007; Monk et al. 2017). To further characterize the impact of the experimental data for BOFdat, the BOFsc of each amino acid were generated from 18 conditions that had both protein weight fractions and proteomic datasets from Schmidt et al., 2016. BOFsc for all samples were then compared to the condition in which cells were grown in presence of glucose, showing that BOFsc indeed varied across conditions (8.74% +/− 2.60%) and that the ones obtained in stationary conditions are the most different (Fig. S3A). We also found that the difference of protein weight fraction from the glucose growth condition was correlated with the average amino acid BOFsc difference (Pearson r = 0.959, p-value = 7.35e-12) indicating that the most important parameter to generate BOFsc is the macromolecular weight fraction.

## 4 Conclusion

BOFdat enables the straightforward determination of BOFsc from mac-romolecular cell composition measurements and omic datasets. Using BOFdat in conjunction with precise inputs obtained from quantitative experimental data standardizes the procedure to generate BOFsc and provides a quality BOF, which is an essential component of GEMs. BOFdat can be used for de novo BOF definition as well as to update pre-existing BOF when more accurate and potentially condition-specific data becomes available.

## Acknowledgements

We thank Jared Broddrick for providing valuable information to determine GAM and NGAM, as well as Nathan Mih for helping with the Pypi distribution.

